# Accelerated somatic mutation calling for whole-genome and whole-exome sequencing data from heterogenous tumor samples

**DOI:** 10.1101/2023.07.04.547569

**Authors:** Shuangxi Ji, Tong Zhu, Ankit Sethia, Wenyi Wang

## Abstract

Accurate detection of somatic mutations in DNA sequencing data is a fundamental prerequisite for cancer research. Previous analytical challenge was overcome by consensus mutation calling from four to five popular callers. This, however, increases the already nontrivial computing time from individual callers. Here, we launch MuSE2.0, powered by multi-step parallelization and efficient memory allocation, to resolve the computing time bottleneck. MuSE2.0 speeds up 50 times than MuSE1.0 and 8-80 times than other popular callers. Our benchmark study suggests combining MuSE2.0 and the recently expedited Strelka2 can achieve high efficiency and accuracy in analyzing large cancer genomic datasets.

## Background

Cancer arises and evolves by accumulating various types of genetic alterations, such as single nucleotide variation (SNV), copy number alteration (CNA) and structural variation (SV). The next-generation sequencing (NGS) technology has revolutionized the way we look at many human diseases, particularly cancer. With its constantly improved capacity and reduced cost, NGS is enabling investigations of genetic alterations within large human patient cohorts, hence advancing both basic and translational cancer research. Many computational tools have been developed for calling somatic variants^1^, which typically require, as input, whole-genome sequencing (WGS) or whole-exome sequencing (WES) data from the tumor tissue, as well as from the blood of the patient to serve as the germline control. WGS provides the most comprehensive coverage to sequence both protein-coding and non-coding regions across the entire genome; whereas WES provides an efficient alternative to WGS by targeting only protein-coding regions that accounts for 1-2% of the genome^2^, hence achieving both higher read depth^3,4^ and lower sequencing cost.

We previously launched MuSE 1.0^5^, a statistical approach for somatic mutation calling, where we introduced a combination of nucleotide base-specific Markov substitution model for molecular evolution and a tumor sample-specific error model to estimate tier-based cutoffs for selecting SNVs. Due to its high sensitivity and specificity, MuSE 1.0 was adopted in multiple pipelines, including as a major contributing caller to reach final consensus calls by the TCGA PanCanAtlas project^6^ across ∼13,000 tumor samples, and the International Cancer Genome Consortium Pan-Cancer Analysis of Whole Genomes (ICGC-PCAWG) initiative^7^ across ∼2,700 tumor samples. One major limitation of MuSE 1.0, like many other mutation callers^8–10^, is the computational speed. It takes 2-3 days to finish running the WGS data of a tumor-normal pair on a typical Linux server with an Intel Xeon processor and more than 100 gigabytes (GB) random access memory (RAM), which explains the commonly seen long wait-times for completing mutation calling before any downstream analysis in large patient cohort studies. Here, we present MuSE 2.0, which maintains the same input, output and mathematical model as MuSE 1.0, but accelerates significantly for both WES and WGS data by adopting a new algorithmic programming backbone. MuSE 2.0 employs a multithreaded producer-consumer model and the OpenMP library for parallel computing, including parsing and uncompressing reads from binary sequence alignment/map formatted (BAM) files, detecting and filtering variants, and writing output. It is also optimized by adopting a more efficient memory allocator. In this paper, we have benchmarked the speed performance of MuSE 2.0 against MuSE 1.0, as well as the other three somatic mutation callers, i.e., MuTect2^9^, SomaticSniper8 and VarScan2^10^, which are the other highlighted somatic mutation callers in the National Cancer Institute Genomic Data Commons (GDC) DNA-seq analysis pipeline (https://docs.gdc.cancer.gov/Data/Bioinformatics_Pipelines/DNA_Seq_Variant_Calling_Pipeline/). To further demonstrate the potential gains, we have benchmarked mutation calling using MuSE2.0 and the other recently expedited mutation caller Strelka2^11^ against consensus mutation calls generated by previous consortial studies using 4 to 5 un-expedited callers^6,7^. We demonstrated the improved utility of our new caller using WES and WGS data generated from 5 tumor-normal pairs with varied average read depths, respectively.

## Results

MuSE 2.0 takes as input the indexed reference genome FASTA file, the BAM format sequencing data from a pair of tumor-normal tissues (**Supplementary Figure 1**) and the dbSNP variant call format (VCF) file, which is bgzip compressed, tabix indexed using the same reference genome. Unlike MuSE 1.0, which can only utilize one core, MuSE 2.0 takes advantage of the multi-core resources in a modern computer or a computing node for somatic SNV calling from WES/WGS data (**Figure 1**).

**Figure 1.**
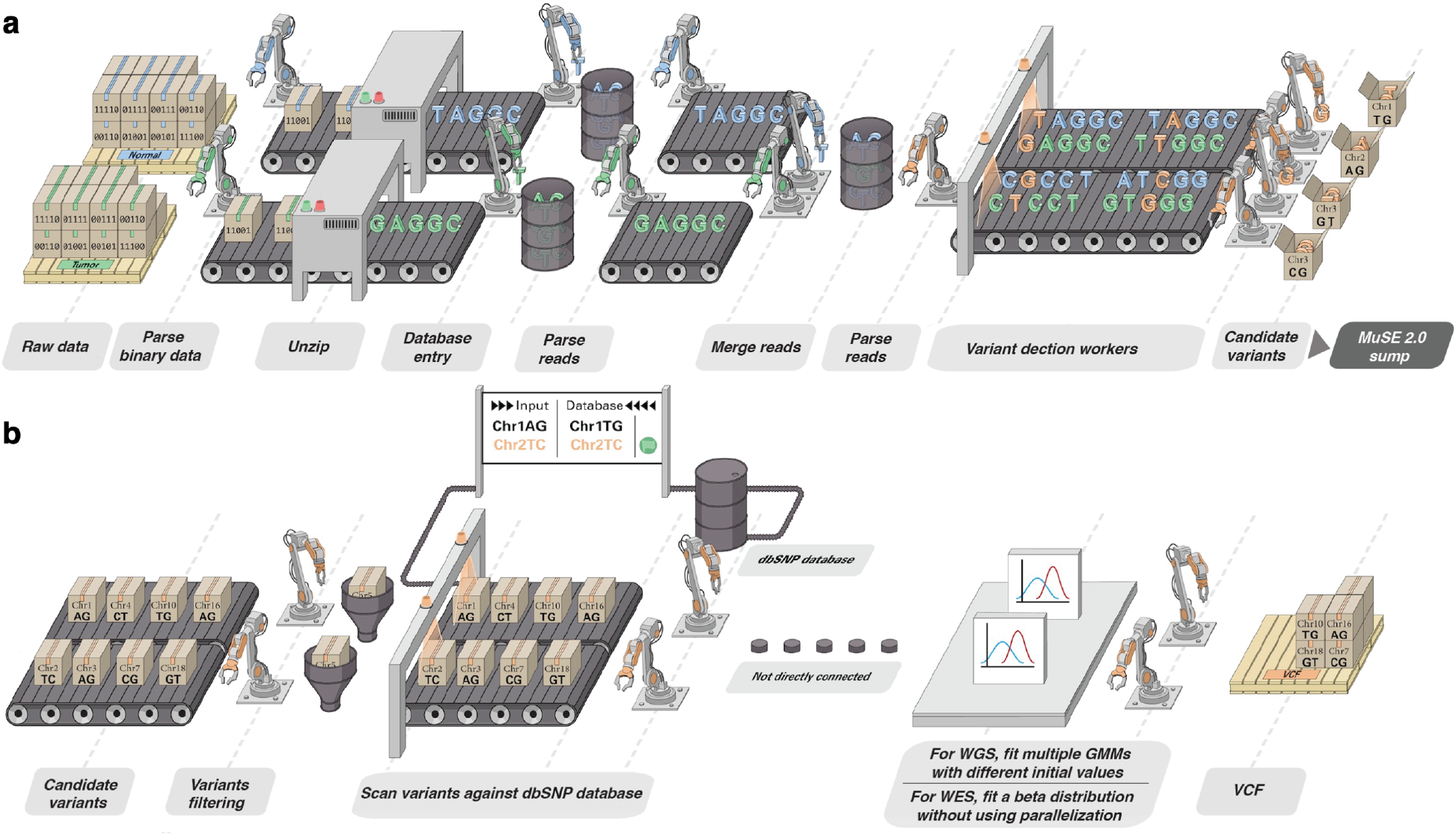
Assembly line illustration of the multi-step parallelization implemented in MuSE 2.0. **a)**, ‘MuSE call’. Workers (threads) keep fetching chunks from the input BAM files from the tumor and normal samples and unzipping them to the text format of reads. Downstream workers combine the reads from tumor and normal samples and send to a queue; from there, other workers detect candidate variants. **b)**, ‘MuSE sump’. Multiple workers are used to take the candidate variants and their corresponding estimated summary statistic π’s and scan them against dbSNP database, labeling those appearing in the database. For candidate variants from the WGS data, we fit two-component Gaussian Mixture Models (GMMs) with multiple initializations, distributed to multiple workers, in order to separate true variants from background noise; for candidate variants from the WES data, no parallelization is implemented due to computational simplicity as we simply fit a beta distribution to π’s.

Since our benchmarking study requires a large amount of computational resources to cover multiple callers and scenarios, we only include results for WES data from 5 tumor-normal pairs and WGS data from 5 tumor-normal pairs, which are randomly selected and downloaded from the Cancer Genome Atlas (TCGA) data portal and the International Cancer Genome Consortium data portal, respectively. The average read coverage of each pre-processed BAM file is show in **Supplementary Table 1**.

We first compare the SNV entries in the output VCF files generated by MuSE 2.0 with those by MuSE 1.0 for each patient sample with the same or different number of cores. Since each SNV entry is denoted by one line of string in a VCF file, we compare the strings from both methods line by line. The result shows that all the entries from the two methods are identical with the same number or different number of cores.

We next compare the speed of running MuSE 2.0 against MuSE 1.0, MuTect2, SomaticSniper, VarScan2 and Strelka2 on a computing cluster. The version information of these methods is listed in **Supplementary Table 2**. Each method is tested with the number of cores at 1, 5, 10, 20, 28, 40 and 80. All methods are assigned with the same RAM of 50GB for WES data and 150GB for WGS data. The time cost of each method for each pair of data is shown in **Figure 2a**. Since MuSE 2.0 and Strelka2 continue to gain computational advantages with increasing number of cores, while the other four methods do not, we examine the overall speed performances of these methods with MuSE 2.0 at 80 cores, Strelka2 at 80 scores, and the average time cost across multiple runs over the different numbers of cores except for core=1 (where the computing resource is too limited) for the other methods (**Figure 2b**). Both MuSE 2.0 and Strelka2 achieve much faster SNV calling compared to all other methods. For WES data, MuSE 2.0 accelerates 28-52 times (mean=44) than MuSE 1.0, 68-83 times (mean=77) than MuTect2, 5-8 times (mean=8) than SomaticSniper, 33-39 times (mean=36) than VarScan2. Similarly, for WGS data, it accelerates 48-59 times (mean=56) than MuSE 1.0, 33-44 times (mean=39) than MuTect2, 7-8 times (mean=8) than SomaticSniper, 33-43 times (mean=36) than VarScan2 for WGS data. Strelka2 achieves about twice the speedup of MuSE 2.0 for both WES and WGS data.

**Figure 2.**
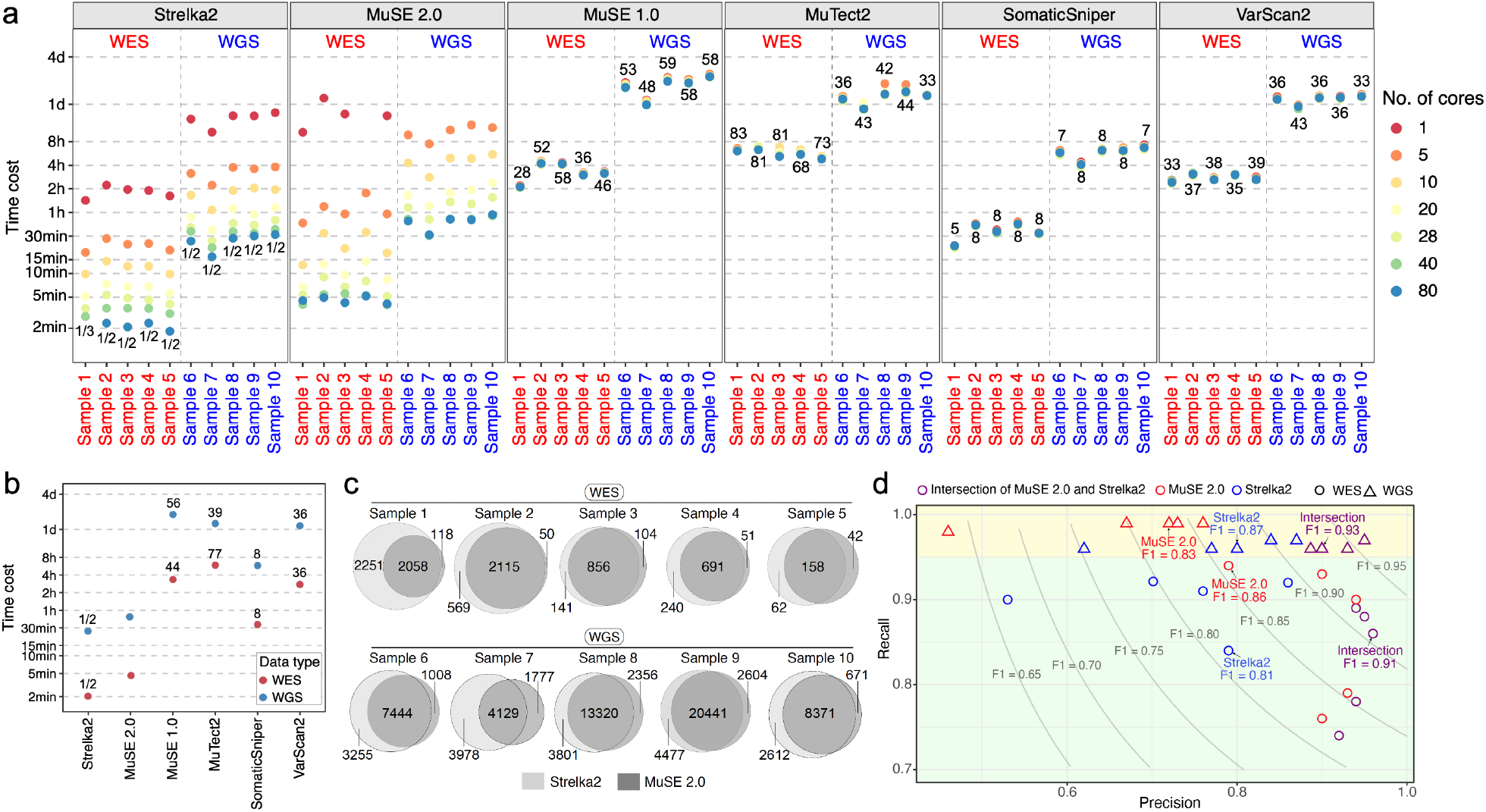
Benchmarking the speed and usability of MuSE 2.0. **a)**, the runtime of MuSE 2.0 against MuSE 1.0 and the other four methods for both WES and WGS data across different numbers of cores. The numbers in the plot represent the fold speedup of MuSE 2.0 (with 80 cores) relative to the other methods whose time cost is averaged across different numbers of cores (excluding No. of core=1). For Strelka2, only the time cost with 80 cores is considered. **b)**, a simplified version of **a)**, in which the time cost of each method is averaged across all samples and different numbers of cores. **c)**, Venn diagrams showing the unique and shared SNV calls of MuSE 2.0 and Strelka2. **d)**, Scatter plot of the precision, recall for the intersection calls from MuSE 2.0 (in red) and Strelka2 (in blue), the calls from each of the two methods (in purple) against the previously reported consensus calls which are considered as the ground truth. For both WES (circle) and WGS (triangle) data, the median F1 scores of the intersection calls, calls from each individual method are shown. Two shaded rectangles highlight the difference of the performance metrics between WES and WGS data. Results from the WGS data are located in the top rectangle.

We further examine the difference between the SNV calls reported by the two expedited methods, MuSE 2.0 and Strelka2, for the same patient sample (**Figure 2c**). For WES data, 46-78% (mean=66%) of the calls are identified by both; 2-16% (mean=7%) of the calls are unique to MuSE 2.0, and 13-51% (mean=27%) of the calls are unique to Strelka2. For WGS data, 42-74% (mean=64%) of the calls are identified by both; 6-18% (mean=11%) of the calls are unique to MuSE 2.0, and 16-40% (mean=25%) of the calls are unique to Strelka2. The difference of the calls motivates us to look into the utility of the intersection calls, and investigate the feasibility of using them to reproduce the consensus calls for these data generated by previous studies^6,7^. These consensus calls previously generated based on 4 to 5 callers are used as the ground truth. We calculate the precision, recall and F1 score, i.e., the harmonic mean of precision and recall, for the intersection calls of each patient sample, as shown in **Figure 2d** and **Supplementary Table 3**. For both WES and WGS data, the intersection calls achieve higher precision values (0.92-0.96 for WES, 0.89-0.95 for WGS), and higher F1 scores (0.82-0.91 for WES, 0.92-0.96 for WGS) compared to the two individual callers. The only exception is that MuSE 2.0 maintains a similar F1 score to the intersection calls with WES data (0.82-0.92). The intersection calls maintain good recall values at 0.74-0.89 (median =0.86) for WES data, and at 0.96-0.97 (median=0.96) for WGS data. The recall values of either the intersection call sets, or the individual call sets from the two methods are consistently higher for WGS data than WES (**Figure 2d**, all results from the WGS data fall in the top rectangle). Also, MuSE 2.0 call sets achieve higher precisions and F1 scores, lower recalls for WES, but higher recalls, lower precisions and F1 scores for WGS data, when compared to the calls from Strelka2. In summary, combining mutation calls from the two expedited callers MuSE 2.0 and Strelka2, e.g., by simply taking an intersection of the calls, is promising to achieve accurate mutation calling in a significantly shorter wait-time, which is particularly useful for WGS data and for analysis of large patient cohorts.

## Discussion

Precision medicine and personalized cancer treatments have advanced remarkably in the last decade, which greatly benefited from the accurate identification of genetic variations in the tumor tissue using NGS data. An efficient and accurate somatic mutation caller is crucial to the scientific studies of all cancers and their clinical management. Previously the accuracy and utility of MuSE 1.0, either alone5 or as a member of a multi-caller consensus calling strategy has been validated by multiple consortial projects^6,7^. This study further develops MuSE 2.0, in order to fully utilize resources on a high-performance computing machine, including both the CPU cores and memory allocation. The producer-consumer model behind the parallelization implemented in the step of ‘MuSE call’ gives MuSE 2.0 the ability to manage multiple processes (workers) at the same time: they run independently at their own rates without being affected by the computing load of other processes. Since the calculation in the step of ‘MuSE sump’ is more straightforward – the computing speed bottlenecks only reside in several for-loop iterations, we therefore use the OpenMP library, with which the parallelization is relatively trivial.

MuSE 2.0 is much faster than the other three benchmarked callers, including MuTect2, SomaticSniper and VarScan2. Although it is slightly slower than Strelka2, its intersection with Strelk2 calls can substantially improve precisions without much loss in recalls, hence improving the overall F1 scores. We therefore demonstrate the utility of the intersection calls from these two fast callers, as compared to using each caller individually or using un-expedited callers. The recalls of the 2-caller intersection calls are lower for WES (TCGA) than WGS (PCAWG). In this case, we suggest adding a third caller such as MuTect2 or more callers, and then take a two out of three consensus approach as previously implemented.

## Conclusion

This study presents MuSE 2.0, improving the mutation calling utility by accelerating its computing speed by up to 50-60 times for both WES and WGS data. MuSE 2.0 reduces the computational time cost of a somatic mutation calling project from ∼40 hours to < 1 hour for WGS data, and from 2-4 hours to ∼5 minutes for WES data, from each pair of tumor-normal samples.

In contrast to the consensus calls from TCGA and PCAWG being generated by five and four callers^6,7^, running MuSE 2.0 and Strelka2 to generate intersection calls may greatly improve the efficiency of genomic data analysis for large patient cohorts, especially for those with WGS data. We therefore expect the proposed MuSE 2.0 to significantly accelerate the variant calling process and benefit the cancer research and clinical communities.

## Methods

### Sample selection

The consensus mutation calls of the TCGA portion of the PCAWG project were downloaded from the ICGC data portal (https://dcc.icgc.org/releases/PCAWG/consensus_snv_indel). The consensus mutation calls of the TCGA MC3 project were downloaded from GDC (https://gdc.cancer.gov/about-data/publications/mc3-2017). We randomly selected 5 patient samples from each of the two repositories, and downloaded the BAM files from the corresponding data portal.

### BAM pre-processing

MuSE 2.0 adopts the same preprocessing steps for the unaligned sequencing reads of the tumor-normal pair as MuSE 1.0, which include trimming poor-quality bases, removing adapters, marking duplicate reads, performing local indel realignment for the paired tumor-normal BAM files jointly, and recalibrating base quality scores (**Supplementary Figure 1**). In this study, the sequencing reads are aligned against the hg19 reference genome build using BWA-MEM12.

### Sequencing depth

The sequencing depth of each BAM file after pre-processing is estimated by samtools with the ‘depth’ command. For WGS data, the overall depth was calculated as the average of the read depths of all genomic locations. For WES data, the overall depth was calculated as the average of the read depths of the genomic locations in the exon regions defined by the exome capture kit downloaded from GDC (https://gdc.cancer.gov/about-data/publications/mc3-2017).

### Parallel computing implementation for MuSE

#### MuSE call

We implement a multithreaded producer-consumer model which deploys threads for parsing and uncompressing reads from BAM files, variant filtering and detection, writing outputs and monitoring the whole process. The model connects all the threads concurrently by thread-safe queues and atomic variables. We also adopt a faster and more efficient memory allocator (i.e., TCMalloc: https://github.com/google/tcmalloc) rather than use the default malloc in C and new in C++ in this step. The parallelization model starts with creating of 6 threads, 3 for the BAM of tumor sample and the other 3 for the BAM of normal sample: 1 of the 3 threads parses the compressed binary data and sends its reference to two queues, namely ChunkReadQueue and ChunkUnzipQueue; the other two threads take the data from the ChunkUnzipQueue, decompress it and label it as ‘processed’. This change is also effective for the data in ChunkReadQueue, since these two queues in fact store the same data. Another thread (i.e., read) is then created, which takes uncompressed data from ChunkReadQueue and recover it to read format for both the BAM tumor sample and the BAM of normal sample, and pushes them to the same queue, ReadQueue. A new thread named processReads is created; it parses the reads from ReadQueue and sends them to the queue, processQ. *n* threads named workers are created to take the reads from processQ and process them following the same pre-filtering and the evolutionary model as MuSE 1.0. The last thread is named as ‘monitor’, which prints the sizes of the queues every second.Here, users can specify *n* according to the number of cores available in the input of MuSE 2.0.

### MuSE sump

We use the OpenMP library to parallelize the three most time-consuming parts in MuSE sump. The first is the loading of candidate variants, the corresponding estimates of equilibrium frequencies for all four alleles (A, C, T, G) for each variant from MuSE call, and filtering out the variants whose ratio between the variant allele frequency from the normal sample and the variant allele frequency from the tumor sample above a predefined cutoff^5^ (0.05). The second is scanning for the remaining variants in the dbSNP and marked as ‘true’ or ‘false’ if they appear in the database or not. For WGS data, MuSE 1.0 fits a two-component Gaussian mixture model to the allele frequencies of the post-filtered variants to separate true mutations from background noise. The parameters (e.g., mean, standard deviation and proportion) of the two components are estimated using the expectation-maximization algorithm which are repeated 50 times with random initializations. For the three parts, we parallel the for-loop iterations using the `‘omp parallel for’` clause from OpenMP in MuSE 2.0 to deploy the computation on multiple cores.

### Speed benchmarking settings

For all the benchmarked methods, if the number of cores requested lies in {1, 5, 10, 20, 28}, the processor is Intel(R) Xeon(R) Gold 6132 CPU @ 2.60GHz; if the number of cores requested lies in {40, 80}, the processor is Intel(R) Xeon(R) Platinum 8380 CPU @ 2.30GHz. We run each method by submitting the LSF (Load Sharing Facility) job script using the bsub command, with which we can easily control the random-access memory (RAM) and the number of cores specified for each method. The options for the 6 callers can be found in **Supplementary Table 4**.

### Precision and recall

We used the consensus SNV calls published previously^6,7^ as the ground truth to evaluate the performance of MuSE 2.0 and Strelka2. For WGS data, we selected all the calls from MuSE 2.0, and only calls from the PASS category from Strelka2 for each patient sample; for WES data, we selected calls from all the categories except for ‘Tier5’ from MuSE 2.0^5^, and only calls from the PASS category from Strelka2 for each patient sample. The intersection of MuSE 2.0 and Strelka2 calls for each patient sample were identified by matching the SNV ids, which combined the columns of CHROM, POS, REF and ALT from the two VCF files. For WES data, we removed the SNVs from the intersection calls outside the regions defined by the exome capture kit of TCGA.

We considered any calls reported by the consensus, but not by the intersection calls as false negatives, any calls reported by the intersection calls, but not by the consensus as false positives. We calculated precision, recall and F1 score to evaluate the accuracy of the new calls.

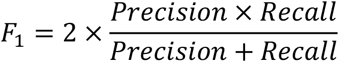

## Availability of data and materials

The ICGC-PCAWG WGS data and the TCGA WES data used in this study can be downloaded from https://dcc.icgc.org/repositories and https://portal.gdc.cancer.gov/repository, respectively. MuSE 2.0 is implemented in C++ and is available at GitHub https://github.com/wwylab/MuSE. A Dockerfile is also included in the repository for building MuSE 2.0 into a Docker image running on Linux machines.

## Acknowledgements

S.J. is supported by the MD Anderson Colorectal Cancer Moon Shot Program and National Institutes of Health (NIH) R01CA268380. W.W. is supported by DoD PC210079, NIH R01CA268380, P30CA016672. S.J. and W.W are also supported by The MD Anderson Cancer Center SPORE in Gastrointestinal Cancer Grant P50 CA221707. We thank Jordan Pietz for his valuable help about making the illustration of Figure 1.

## Author contributions

S.J. implemented the parallelization for the ‘MuSE sump’ step, performed the speed benchmarking, analyzed the results, and wrote the manuscript in collaboration with other authors.T.Z. and A.S. implemented the parallelization for the ‘MuSE call’ step. W.W. conceived the project, planned, and supervised the work, wrote the manuscript, in collaboration with all other authors. All authors commented on and approved the final manuscript.

## Supplementary information

**Supplementary Figure 1.**
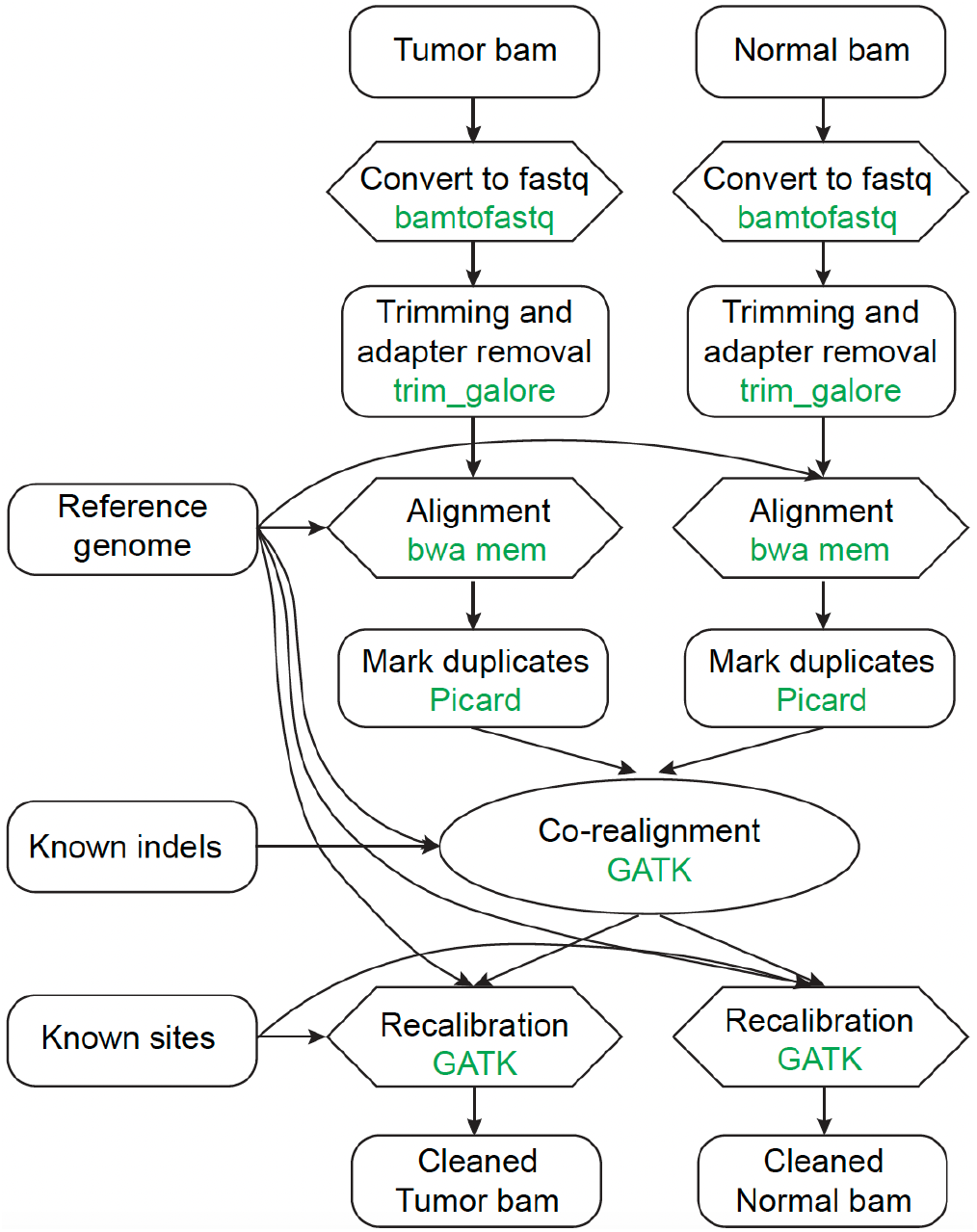
Flowchart of sequencing read pre-processing followed by all methods in the benchmarking study.

**Supplementary Table 1.**
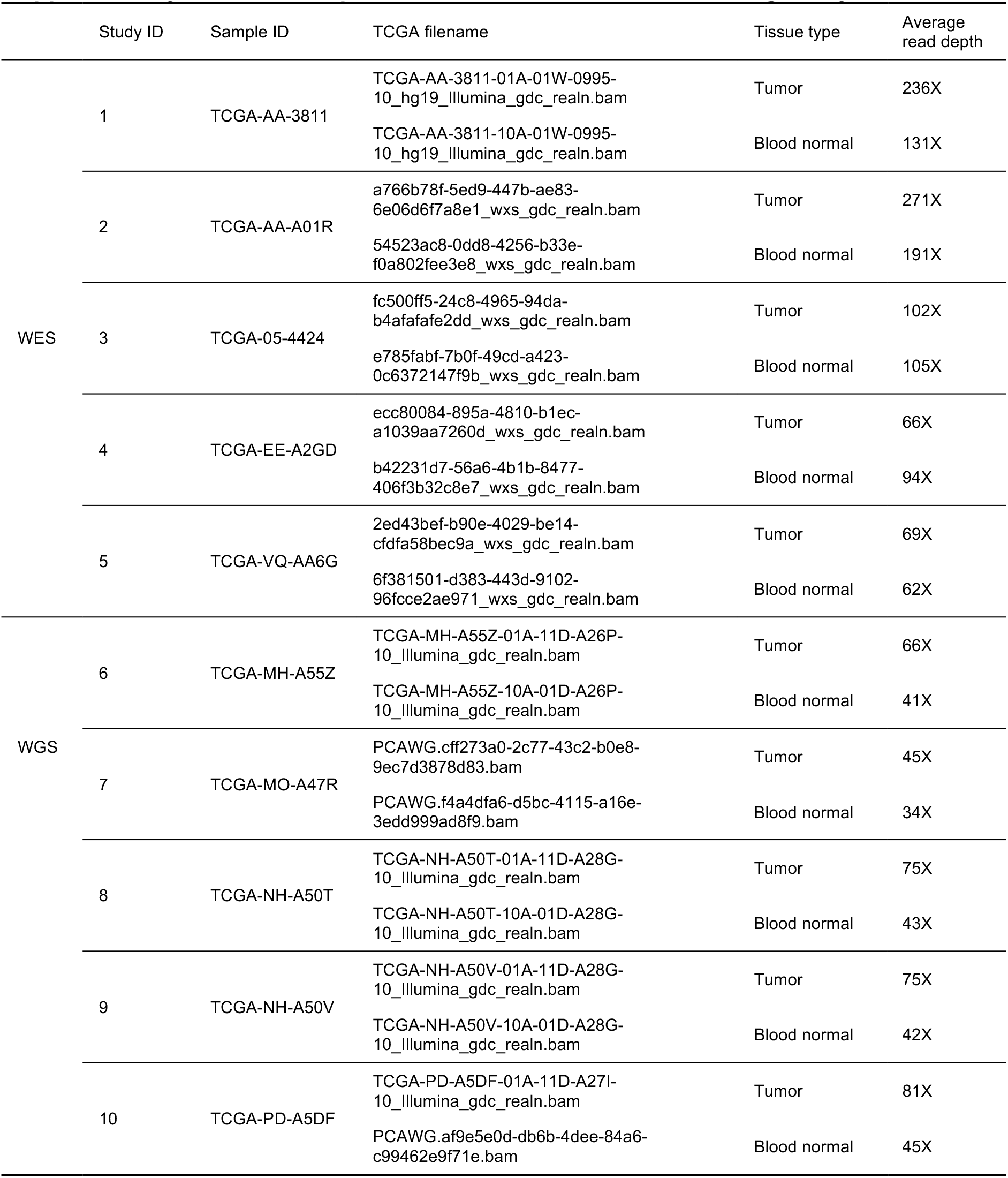
Sample information for the benchmarking study.

**Supplementary Table 2.** Version information of the benchmarked tools.

| Tool | Version |
| --- | --- |
| MuSE 2.0 | 2.0.1 |
| MuSE 1.0 | 1.0rc |
| MuTect2 | 4.1.9.0 |
| SomaticSniper | 1.0.5.0 |
| Varscan2 | 2.4.2 |
| strelka2 | 2.9.x |

**Supplementary Table 3.** Precision, recall and F1 scores of the calls from Strelka2, MuSE 2.0 and their intersections for both WES and WGS data.

| Sample ID | Data type | Intersection of MuSE 2.0 and Strelka2 |  |  | Strelka2 |  |  | MuSE 2.0 |  |  |
| --- | --- | --- | --- | --- | --- | --- | --- | --- | --- | --- |
|  |  | Precision | Recall | F1 score | Precision | Recall | F1 score | Precision | Recall | F1 score |
| 1 | WES | 0.92 | 0.74 | 0.82 | 0.53 | 0.90 | 0.67 | 0.90 | 0.76 | 0.82 |
| 2 |  | 0.94 | 0.78 | 0.85 | 0.79 | 0.84 | 0.81 | 0.93 | 0.79 | 0.85 |
| 3 |  | 0.95 | 0.88 | 0.91 | 0.86 | 0.92 | 0.89 | 0.90 | 0.93 | 0.91 |
| 4 |  | 0.96 | 0.86 | 0.91 | 0.76 | 0.91 | 0.83 | 0.94 | 0.90 | 0.92 |
| 5 |  | 0.94 | 0.89 | 0.91 | 0.70 | 0.92 | 0.80 | 0.79 | 0.94 | 0.86 |
| 6 | WGS | 0.95 | 0.97 | 0.96 | 0.84 | 0.97 | 0.90 | 0.67 | 0.99 | 0.80 |
| 7 |  | 0.89 | 0.96 | 0.92 | 0.62 | 0.96 | 0.76 | 0.46 | 0.98 | 0.63 |
| 8 |  | 0.90 | 0.96 | 0.93 | 0.77 | 0.96 | 0.86 | 0.72 | 0.99 | 0.83 |
| 9 |  | 0.89 | 0.96 | 0.92 | 0.8 | 0.96 | 0.87 | 0.76 | 0.99 | 0.86 |
| 10 |  | 0.93 | 0.96 | 0.94 | 0.87 | 0.97 | 0.91 | 0.73 | 0.99 | 0.84 |

**Supplementary Table 4.** Running commands of the benchmarked tools.

| Tool | Command |
| --- | --- |
| MuSE 2.0 | `MuSE call -O muse -f -n 10 $reference tumor.bam normal.bam MuSE sump -I muse.MuSE.txt -O muse.vcf -n 20 -E -D dbsnp` |
| MuSE 1.0 | `MuSE call -O muse -f $reference tumor.bam normal.bam MuSE sump -I muse.MuSE.txt -O muse.vcf -E -D dbsnp` |
| MuTect2 | `gatk Mutect2 -R reference_genome -I tumor.bam -I normal.bam -tumor tumor_id -normal normal_id -O mutect2_raw.vcf.gz gatk FilterMutectCalls -V mutect2_raw.vcf.gz -R reference_genome -O mutect2_filtered.vcf.gz` |
| **SomaticSniper** | `bam-somaticsniper -q 1 -L -G -Q 15 -s 0.01 -T 0.85 - N 2 -r 0.001 -n NORMAL -t TUMOR -F vcf -f reference_genome tumor.bam normal.bam somaticsniper.vcf` |
| **Varscan2** | `samtools mpileup -f $reference -q 1 -B normal.bam tumor.bam > mpileup.pileup java -jar VarScan.v2.4.1.jar somatic mpileup.pileup varscan_somatic.vcf --mpileup 1 --min-coverage 8 -- min-coverage-normal 8 --min-coverage-tumor 6 --min- var-freq 0.10 --min-freq-for-hom 0.75 --normal- purity 1.0 Competing interest statement --tumor- purity 1.00 --p-value 0.99 --somatic-p-value 0.05 -- strand-filter 0 --output-vcf java -jar VarScan.v2.4.1.jar processSomatic varscan.vcf.snp --min-tumor-freq 0.10 --max-normal- freq 0.05 --p-value 0.07` |
| **strelka2** | `STRELKA_INSTALL_PATH= ${STRELKA_INSTALL_PATH}/bin/configureStrelkaSomatic Workflow.py --normalBam normal.bam --tumorBam tumor.bam --referenceFasta reference_genome -- runDir ./ ./runWorkflow.py -m local -j 20` |

